# Noninvasive Prenatal Diagnosis for Duchenne Muscular Dystrophy Based on the Direct Haplotype Phasing

**DOI:** 10.1101/720987

**Authors:** Min Chen, Chao Chen, Xiaoyan Huang, Jun Sun, Lu Jiang, Yingting Li, Yaping Zhu, Changgeng Tian, Yufan Li, Zhe Lu, Yaoshen Wang, Fanwei Zeng, Yun Yang, Zhiyu Peng, Chenghong Yin, Dunjin Chen

**Affiliations:** Department of Fetal Medicine and Prenatal Diagnosis, the Third Affiliated Hospital of Guangzhou Medical University, Guangzhou, 510150, China; Obstetrics & Gynecology Institute of Guangzhou, Guangzhou, 510150, China; The Medical Centre for Critical Pregnant Women in Guangzhou, Guangzhou, 510150, China; Key Laboratory for Major Obstetric Diseases of Guangdong Province, Guangzhou, 510150, China; Tianjin Medical Laboratory, BGI-Tianjin, BGI-Shenzhen, Tianjin 300308, China; BGI Genomics, BGI-Shenzhen, Shenzhen, 518083, China; Wuhan BGI Clinical Laboratory Co., Ltd, BGI-Wuhan, BGI-Shenzhen, Wuhan 430074, China; Beijing Obstetrics and Gynecology Hospital, Capital Medical University, Beijing 100026, China; Key Laboratory for Reproduction and Genetics of Guangdong Higher Education Institutes

## Abstract

**Objective:** We aimed to investigate the validity of noninvasive prenatal diagnosis (NIPD) based on direct haplotype phasing without the proband and its feasibility for clinical application in the case of Duchenne Muscular Dystrophy (DMD).

**Methods:** Thirteen singleton-pregnancy families affected by DMD were recruited. Firstly, we resolved maternal haplotypes for each family by performing targeted linked-read sequencing of their high molecular weight DNA, respectively. Then, we identified SNPs of the *DMD* gene in all carrier mothers and inferred the *DMD* gene mutation status of all fetuses. Finally, the fetal genotypes were further validated by using chorionic villus sampling.

**Results:** The method of directly resolving maternal haplotype through targeted linked-read sequencing was smoothly performed in all participated families. The predicted mutational status of 13 fetuses was correct, which had been confirmed by invasive prenatal diagnosis.

**Conclusion:** Direct haplotyping of NIPD based on linked-read sequencing for DMD is accurate.

## Introduction

Duchenne Muscular Dystrophy (DMD) is an X-linked recessive disease characterized by progressive muscle degeneration and weakness^1^, which usually occurs in early childhood, affecting approximately 1 in 3500 male births worldwide^2^. It is reported that DMD is caused by the mutations of the *DMD* gene on the X chromosome^3^. Current gold standards for prenatal diagnostic testing include amniocentesis or chorionic villus sampling (CVS)^4^. With the discovery of cell-free fetal DNA in maternal plasma^5^ and the development of genetic testing methods, noninvasive prenatal diagnosis (NIPD) is increasingly used in clinical practice. The feasibility and accuracy of detecting fetal DNA using maternal plasma DNA have been confirmed by a large number of studies. NIPD was initially used to detect fetal aneuploidy, but it’s application in single gene disorders has also been explored^6^.

The detection of DMD by NIPD has been researched previously, but most studies are based on trios’ strategy with a proband^7,8^. This approach can address some of the needs of prenatal diagnosis, but for female carriers, it is unable to detect the genotypes of their first child because there is no proband could provide assistance for resolving haplotypes. Many efforts have been made to overcome this problem, with microfluidics-based linked-read sequencing technology being used prominently^9^. This is a direct haplotyping phasing approach and has been reported to predict fetal genotypes without a proband in a family accurately. In the study published by Hui et al.^10^, the mutation inheritance status of the fetus was successfully deduced by integrating the linked-read sequencing technology of 10X Genomics, next-generation targeted sequencing and relative haplotype dosage (RHDO) analysis^11^. To confirm the accuracy of the direct haplotype-based NIPD and its feasibility for clinical application in the case of DMD, we recruited 13 families affected by DMD for research.

## Methods

### Sample collection

We recruited 13 high-risk families without a proband at the Third Affiliated Hospital of Guangzhou Medical University and obtained informed consent. All procedures were performed following the tenets of the Declaration of Helsinki and were approved by the Ethics Committee of the hospital. For each family, we collected 10ml maternal blood samples and 5mg CV between 11-25 weeks.

### Maternal DNA linked-read sequencing

In this study, we designed a 657.29Kb SeqCap kit (Roche, Basel, Switzerland) for the *DMD* gene detection, which contains the coding region (13.91Kb) and the flanking region (1M) of the *DMD* gene, and the gender determination locus on the Y chromosome. The peripheral blood samples of the 13 carrier mothers were collected to obtain the genomic DNA (gDNA). Then, we isolated maternal high-molecular weight gDNA (> 40kb) to get barcoded DNA fragments by processing with 10X Genomics ChromiumTM library (Pleasanton, CA) protocol. Finally, we performed targeted linked-read sequencing by using PE101 bp on Illumina Hiseq2500 sequencing system (San Diego, CA).

### Plasma DNA sequencing library preparation

We used the QIAamp Circulating Nucleic Acid kit to extract cell-free DNA from maternal plasma. Then, we prepared the library according to the protocol and performed sequencing on the Illumina platform (Hiseq 2500).

### Sequence alignment and SNP calling

We aligned the barcoded reads of maternal to the reference genome (GRCh37/hg19) by performing 10X Lariat™ and called variants by using the GATK method in Long Ranger. The paired-end sequencing reads of maternal plasma DNA were mapped to the reference genome (GRCh37/hg19) using SOAP2 and the variants were determined using GATK software.

### Haplotype phasing

We resolved the maternal haplotypes from linked-read sequencing data of gDNA. Sequenced reads that shared the same barcode with the one carrying mutant allele were linked (mutant-linked barcode reads) and phased to the same haplotype. Wild-type linked barcode reads were phased to the opposite haplotype. Heterozygous SNPs associated with same haplotypes were used for subsequent fetal genotypic prediction and recombinant detection.

### Direct haplotype-based NIPD of the fetus

Because DMD is X chromosome inheritance, we focused on the SNPs of the X chromosome in the following analysis process. We constructed the Hidden Markov Model (HMM) by using maternal heterozygous SNPs on chromosome X and maternal plasma sequencing data to infer fetal haplotype. For each site, the HMM emission probabilities were the probabilities of genetically pathogenic and non-pathogenic alleles in the fetus, which were calculated by analyzing the number of reads in maternal plasma. The HMM transition probabilities were the recombination rates between SNPs, which were obtained from the SNPs genetic map (from NCBI). When analyzing the HMM model, we used the Viterbi algorithm to deduce the inherited haplotype and recombination breakpoints in the fetus.

### Validation of NIPD for DMD

To further verify the accuracy of NIPD, we obtained the fetal DNA by chorionic villus sampling and performed targeted sequencing using the same probe.

## Results

Thirteen families at risk for a fetus with DMD were recruited. The clinical information and the mutational status of the studied cases are listed in Table 1.

**Table 1.** Clinical Information and Molecular Diagnosis.

| Family | Sample | GW | DMD mutation |
| --- | --- | --- | --- |
| F01 | mat |  | Het.D.EX45-48 |
|  | plasma | 19W+1D | - |
| F02 | mat |  | Het.D.EX49-52 |
|  | plasma | 12W+1D | - |
| F03 | mat |  | Het.D.EX48-49 |
|  | plasma | 11W+2D | - |
| F04 | mat |  | Het.D.EX46 |
|  | plasma | 11W+5D |  |
| F05 | mat |  | Het.c.1525TC[2] |
|  | plasma | 12W+6D |  |
| F06 | mat |  | Het.D.EX50 |
|  | plasma | 12W+1D |  |
| F07 | mat |  | Het.D.EX45 |
|  | plasma | 17W |  |
| F08 | mat |  | Het.D.EX45 |
|  | plasma | 12W+4D |  |
| F09 | mat |  | Het.c.237_256delGGCACTGCGGGTTTTGCAGA |
|  | plasma | 17W+3D |  |
| F10 | mat |  | Het.c.10141C>T |
|  | plasma | 11W+3D |  |
| F11 | mat |  | Het.D.EX42-43 |
|  | plasma | 13W+3D |  |
| F12 | mat |  | Het.D.EX3-13 |
|  | plasma | 12W+4D |  |
| F13 | mat |  | Het.c.1408A > T |
|  | plasma | 25W+4D |  |

Notes: Abbreviations: mat, maternal; Het. D.EX, Heterozygous deletion Exon; GW, Gestational Weeks.

The results of the targeted linked-read sequencing of all the samples were listed in Table 2. The mean depth of 13 maternal gDNA samples is 561X (rang: 329X-697X) and the average length of N50 phase-block is 741.61kb (range from 341.37kb to 963.80kb). Through the bioinformatics analysis, there were more than 98% reads would map to the reference genome (GRCH37/hg19).

**Table 2.** Statistics of 10X Linked-read Sequencing Data.

| **Family** | **Gems detected** | **SNPs phased** | **Longest phase block** | **N50 phase block** | **Mean depth of target region** | **Mapped reads** |
| --- | --- | --- | --- | --- | --- | --- |
| F01 | 377998 | 97.95% | 2993951 | 798365.00 | 329 | 98.95% |
| F02 | 482807 | 98.20% | 3110821 | 963803.00 | 531 | 98.94% |
| F03 | 488937 | 98.05% | 2474849 | 963803.00 | 537 | 99.09% |
| F04 | 572612 | 98.35% | 1366582 | 479287.00 | 697 | 99.48% |
| F05 | 549633 | 98.26% | 1983343 | 493754.00 | 644 | 99.58% |
| F06 | 568884 | 98.08% | 1376475 | 341366.00 | 664 | 99.60% |
| F07 | 457130 | 98.37% | 2437345 | 825353.00 | 653 | 99.42% |
| F08 | 406176 | 98.06% | 1806841 | 579443.00 | 532 | 99.57% |
| F09 | 494357 | 98.31% | 1950712 | 543877.00 | 584 | 99.61% |
| F10 | 506946 | 98.58% | 1799819 | 612603.00 | 637 | 99.38% |
| F11 | 431524 | 98.35% | 1873856 | 562824.00 | 581 | 99.53% |
| F12 | 424937 | 98.21% | 3494973 | 1498192.00 | 508 | 98.96% |
| F13 | 392900 | 98.11% | 4084221 | 978321.00 | 405 | 98.99% |

Reads that shared the same barcode and had the mutant allele at heterozygous SNP positions were considered to be the mutant-linked reads, designated as Hap0, while reads with the wild-type allele at same SNP positions were termed as Hap1. We directly resolved the 2 haplotypes of all 13 maternal gDNA by linking the haplotype blocks assembled by the barcoded reads.

The plasma DNA samples from 13 carrier mothers were target sequenced in the DMD region. The fraction of cffDNA ranged from 7.45% to 15.39% (Table 3). We used informative SNPs (average 701, Table 3) to construct the fetal haplotypes and performed HMM to infer the genotype of fetal *DMD*. The results of NIPD showed that four fetuses had inherited the Hap0 maternal haplotype, including two female carriers (F01and F08) and two affected male fetuses (F02 and F07), and that seven fetuses had inherited the Hap1 maternal haplotype, including six normal female fetuses (F03, F04, F06, F09, F10, F11and F12) and two normal male fetuses (F05 and F13) (Table 3 and Figure 1).

**Table 3.** Noninvasive Prenatal Diagnosis of DMD Families.

| Family | Fetal DNA fraction | Fetal gender | Phased SNPs | Hap0 | Hap1 | Fetal Haplotype | NIPD | CVS |
| --- | --- | --- | --- | --- | --- | --- | --- | --- |
| F01 | 15.39% | female | 790 | 606 | 184 | Hap0 | carrier | carrier |
| F02 | 6.22% | male | 877 | 877 | 0 | Hap0 | affected | affected |
| F03 | 10.13% | female | 774 | 0 | 774 | Hap1 | normal | normal |
| F04 | 8.60% | female | 435 | 0 | 435 | Hap1 | normal | normal |
| F05 | 8.60% | male | 781 | 0 | 781 | Hap1 | normal | normal |
| F06 | 12.32% | female | 262 | 0 | 262 | Hap1 | normal | normal |
| F07 | 9.14% | male | 727 | 718 | 9 | Hap0 | affected | affected |
| F08 | 18.54% | female | 750 | 750 | 0 | Hap0 | carrier | carrier |
| F09 | 7.45% | female | 860 | 0 | 860 | Hap1 | normal | normal |
| F10 | 7.68% | female | 384 | 0 | 384 | Hap1 | normal | normal |
| F11 | 9.67% | female | 425 | 0 | 425 | Hap1 | normal | normal |
| F12 | 8.21% | female | 964 | 0 | 964 | Hap1 | normal | normal |
| F13 | 16.97% | male | 1087 | 0 | 1087 | Hap1 | normal | normal |

Notes: Hap0, fetal-inherited maternal mutant haplotype; Hap1, fetal-inherited maternal wild-type
haplotype; CVS, Chorionic Villi Sampling ;

**Figure 1.**
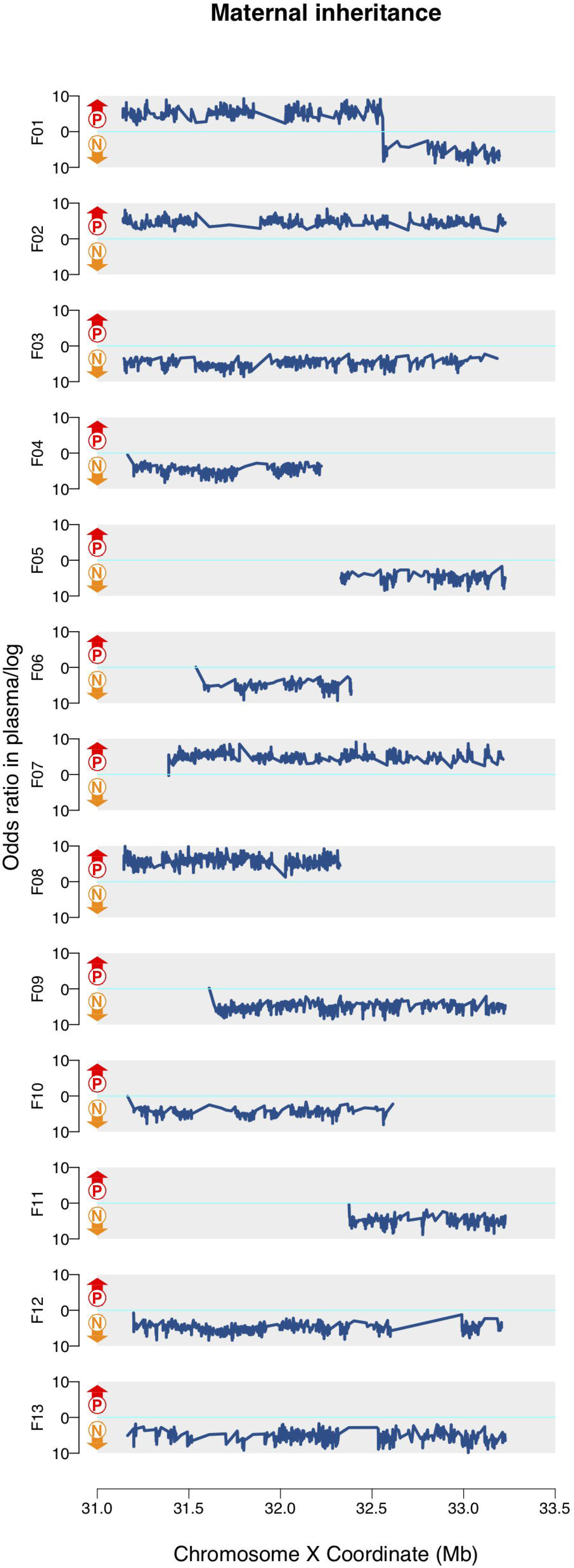
Fetal haplotype determination. The blue lines represent the maternal haplotype of fetal inheritance on the X chromosome. The lines above zero (Cyan line) indicate that the pathogenic allele is inherited (Hap0), while the lines below zero mean the normal allele is inherited (Hap1)..

To further evaluate the performance of the direct haplotype-based NIPD, we performed targeted capture sequencing in CV samples, and all families had the same results as NIPD (Table 3).

## Discussion

Traditional prenatal diagnostic methods, including chorionic villus sampling, amniocentesis, and umbilical cord puncture can be invasive and may put both the fetus and the pregnant woman at risk. The NIPD method is simpler and safer than invasive diagnosis because it only requires blood testing. Since the introduction of NIPD to clinical practice in 2011^12^, it has been widely implemented and is currently offered in over 60 countries^13^. NIPD was initially used to test for Down’s Syndrome and other forms of fetal aneuploidy by using maternal plasma cell-free fetal DNA^14^. Now, it is also used for testing for monogenic diseases^6^. However, the limitations of NIPD applications in determining monogenic disorder are becoming more prominent as it requires genomic data from parents and the proband to resolve the parental haplotypes.

To the best of our knowledge, two methods have recently been proposed to overcome this disadvantage. Hui et al. have successfully applied linked-read sequencing technology in combination with relative haplotype dosage analysis to noninvasive prenatal testing of families without proband^10^. Vermeulen et al. combined the targeted locus amplification (TLA)-based haplotype phasing of the two parents with targeted sequencing of cell-free DNA for noninvasive prenatal testing of families without a first affected child^15^. These two-direct haplotype-based NIPD methods could accurately deduce the fetal mutation status without relying on the availability of DNA from the proband. In comparison, targeted haplotyping is less expensive, and linked-read sequencing has an advantage in recombination events analysis. The application of the two approaches in the case of DMD requires further exploration.

In this study, we first directly resolved the maternal haplotypes by performing linked-read sequencing, and then successfully predicted the genotype of fetal *DMD* by using the NIPD method in all 13 families. Gender identification is essential for the NIPD of X-linked recessive genetic diseases. We determine the sex of the fetus based on the average depth and the coverage of the target region in chromosome Y in the plasma samples, as described in our previous work^16^.

Due to the limited researches in this area to date, more studies are needed to prove its clinical feasibility. Our study demonstrated that the direct haplotype-based method could provide accurate fetal genotype information on DMD without the proband, meaning that it could be an additional option for the NIPD of DMD.

## Conclusion

The direct haplotype-based approach for DMD has potential for introduction for clinical service. To the best of our knowledge, this is the largest cohort without the probands for NIPD of DMD. Further studies should aim to refine the methodology, reduce the cost, and adopt this approach for a larger population with a wide range of genetic disorders.

## Acknowledgments

We thank all participants in this study for their collaborations.

The work was supported by the National Key Research and Development Program of China (2018YFC1004104),the National Natural Science Foundation of China (NSFC) (No. 81671470),, Guangzhou Science and Technology Program (No. 2014 Y2-00551, No. 201504282321393, No. 201604020078, No.201604020091), Guangdong Science and Technology Program (No. 2013B022000005, No. 2016A030313610).

